# The ENL–USP7 Complex Regulates HIV Latency Through BRD4 Stabilization

**DOI:** 10.64898/2026.04.08.717209

**Authors:** Waqas Ahmed, Xiaoyi Li, Ciniso S. Shabangu, Hongjie Chen, Nikesh Kutwal, Jennifer Kirchherr, Wenshe R. Liu, Xin Li, Yongcheng Song, Kaixiu Luo, H. Umit Kaniskan, Jian Jin, Sara Gianella, David M. Margolis, Nancie M. Archin, Yuyang Tang, Guochun Jiang

**Affiliations:** UNC HIV Cure Center and Institute of Global Health and Infectious Diseases, The University of North Carolina at Chapel Hill, Chapel Hill, NC 27599-7042, USA; Department of Chemistry, Texas A&M University, College Station, TX, 2007, USA; Verna and Marrs McLean Department of Biochemistry and Molecular Pharmacology, Baylor College of Medicine, 1 Baylor Plaza, Houston, TX 77030, USA; Mount Sinai Center for Therapeutics Discovery, Departments of Pharmacological Science, Oncological Science, and Neuroscience, The Mount Sinai Tisch Cancer Center, Icahn School of Medicine at Mount Sinai, New York, NY 10029, USA; Department of Medicine, University of California at San Diego, La Jolla, CA, 92093 USA; UNC HIV Cure Center, Institute of Global Health and Infectious Diseases, and Department of Biochemistry and Biophysics, The University of North Carolina at Chapel Hill, Chapel Hill, NC 27599, USA

**Keywords:** HIV latency, HIV reservoir, crotonylation, ENL, USP7, BRD4, SEC, microglia

## Abstract

HIV-1 persists in CD4⁺ T cells and brain microglia through host factors that enforce viral latency, yet the mechanisms that stabilize key transcriptional regulators remain incompletely understood. Here, we identify the YEATS domain-containing protein ENL and its associated deubiquitinase USP7 as a host complex that maintains HIV-1 latency. USP7 stabilizes BRD4 by deubiquitination, suppressing HIV transcription and sustaining viral quiescence. Disruption of the ENL–USP7 complex using selective PROTACs reactivates latent HIV in cell line models, as well as in resting CD4⁺ T cells and microglia isolated from people with HIV on antiretroviral therapy. These findings uncover a critical ENL–USP7–BRD4 axis that enforces HIV-1 latency and highlight USP7 as a potential target for latency-reversing strategies.

**Highlights:**

- ENL, a YEATS domain-containing crotonylation reader, acts as a suppressor rather than an activator of HIV-1 transcription.
- ENL recruits USP7 to stabilize BRD4 and enforce viral latency.
- Disruption of the ENL–USP7–BRD4 axis reactivates latent HIV in T cells and microglia.
- Targeting USP7 or ENL reveals a therapeutic vulnerability in HIV reservoirs.

## Introduction

Efforts to cure HIV-1 are hindered by the persistence of long-lived infected immune cells that harbor transcriptionally silent proviruses and evade immune clearance, underscoring the need to define the host and viral mechanisms that sustain HIV-1 latency ^1,2^. Although antiretroviral therapy (ART) effectively suppresses viral replication, it does not eliminate the latent reservoir, allowing intermittent proviral reactivation and rapid viral rebound upon treatment interruption, necessitating lifelong therapy ^3,4^. HIV-1 latency is maintained by multiple barriers operating at epigenetic, transcriptional, and post-transcriptional levels ^5^.

A key regulatory checkpoint in HIV transcription is the transition of RNA polymerase II into productive elongation, a process controlled by host elongation factors assembled within the super elongation complex (SEC), which has been widely proposed as a critical activator of HIV-1 transcription ^6^. The SEC includes components such as P-TEFb, ELL2, AFF1/4, and the YEATS domain-containing protein ENL (MLLT1), which functions as a reader of histone acylation marks to facilitate chromatin recruitment and release of promoter-proximal paused RNA polymerase II ^7–10^.

Unexpectedly, emerging evidence challenges this model, suggesting that the SEC may instead repress HIV transcription at the viral promoter ^11–14^. However, the molecular basis for this unexpected activity remains unclear.

Here, we identify the acylation reader ENL as an unexpected regulator of HIV-1 latency that functions as a transcriptional suppressor rather than an activator. Mechanistically, ENL cooperates with the deubiquitinase USP7 to stabilize the latency-promoting factor BRD4, thereby enforcing proviral quiescence. Disruption of this ENL–USP7–BRD4 axis destabilizes BRD4 and reactivates latent HIV. These findings reveal a distinct SEC-associated mechanism that promotes, rather than relieves, HIV transcriptional silencing.

## Results

### ENL functions as an important regulator of HIV transcription and latency

ENL, a core subunit of the SEC, plays a key role in regulating transcriptional elongation in many host genes ^7,15^. It has been proposed that ENL is within the SEC at the HIV LTR, thus contributing to HIV transcription ^9^. However, direct evidence remains lacking. To assess the effect of ENL inhibition on HIV-1 latency, Jurkat cell-derived HIV latency models (2D10, J-Lat A1) were treated with SR-0813, a selective ENL/AF9 YEATS domain inhibitor ^16^, for 24 h. Treatment with SR-0813 increased HIV-driven GFP expression in 2D10 and J-Lat A1 cells (**Fig. 1A, B)**, as measured by flow cytometry, without detectable cellular toxicity (**Fig. S1A, B**), and also induced HIV transcription from latency in 2D10 and J-Lat A1 cells **(Fig. 1C)**. Furthermore, another recently discovered ENL-specific YEATS-domain inhibitor, ENLi13 ^17^, also markedly reactivated latent HIV-1 proviruses, as evidenced by increased GFP expression **(Fig. 1D, E)** and increased HIV transcription in both HIV latency models **(Fig. 1F)**, without detectable cellular toxicity **(Fig. S1C, D)**.

**Figure 1.**
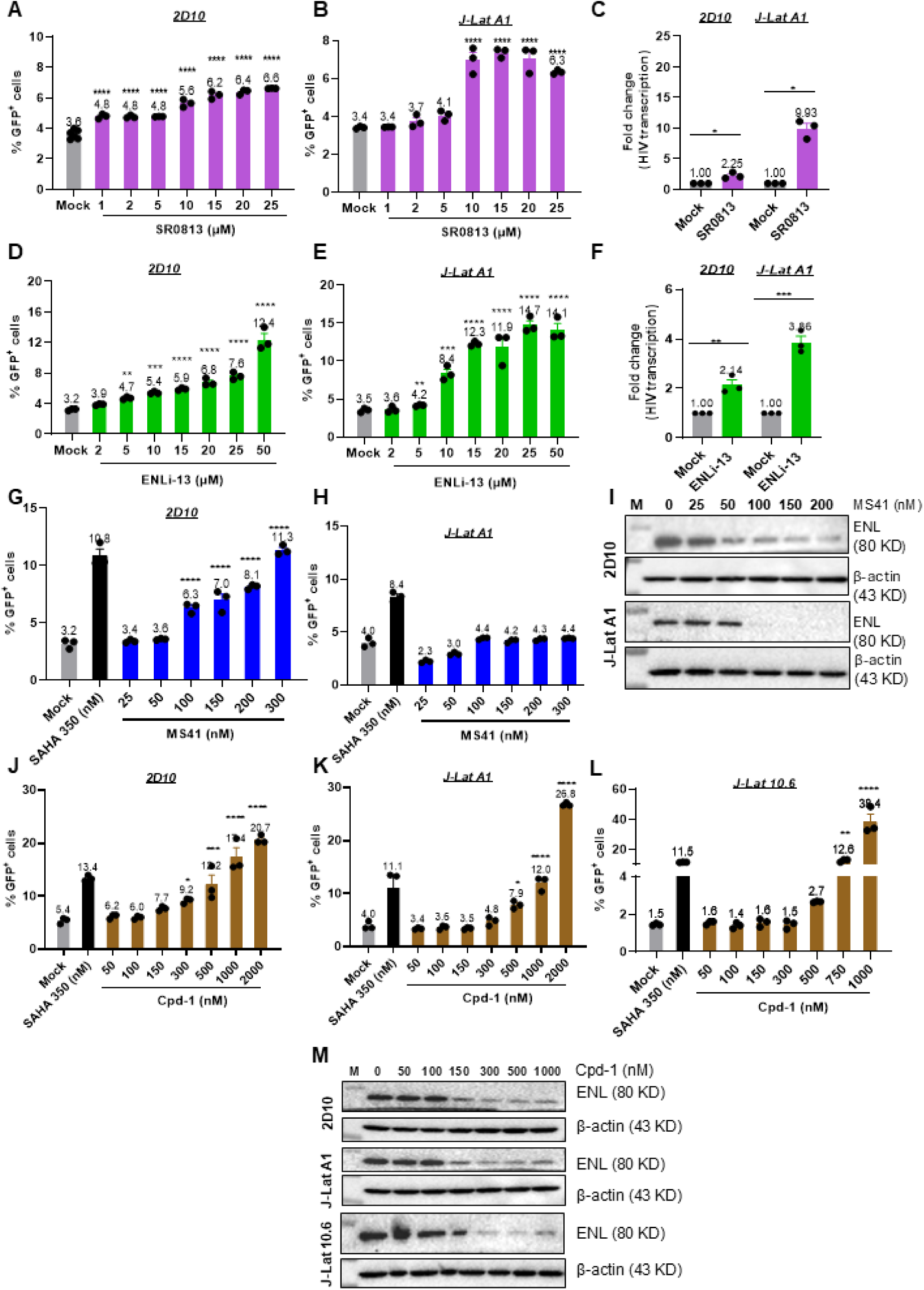
ENL inhibition or protein degradation induces HIV transcription from latency in HIV latency models. Jurkat latency models (2D10, J-Lat A1, and J-Lat 10.6) were treated with increasing concentrations of the ENL inhibitors SR0813 or ENLi-13, or the ENL-targeting PROTACs MS41 and Cpd-1. Cells were harvested 24 h post-treatment, and HIV reactivation was quantified by GFP expression using flow cytometry, while HIV transcription was assessed by qPCR (n = 3). Dose-dependent GFP induction in 2D10 and J-Lat A1 cells following treatment with SR0813 (A and B), ENLi-13 (D and E), MS41 (G and H), and Cpd-1 (J and K) as well as in J-Lat 10.6 cells (L) is shown. HIV transcription in 2D10 and J-Lat A1 cells upon SR0813 (C) and ENLi-13 (F) is shown. The *P* values were determined using one-way ANOVA or two-tailed Student’s t-test. Dose-dependent ENL degradation assessed by immunoblotting in 2D10 and J-Lat A1 cells treated with DMSO or MS41 (I), or Cpd-1 for 24 h (M), with β-Actin as a loading control.

The ENL-targeting proteolysis-inducing chimera (PROTAC) MS41 ^18^, a potent and selective VHL-recruiting ENL degrader, robustly reactivated HIV in 2D10 cells when applied at 100-300 nM for 24 h **(Fig. 1G; Fig. S1E)** but had no detectable effect in J-Lat A1 cells **(Fig. 1H; Fig. S1F)**. Consistent with the previous report in cancer cells, MS41 also markedly induced ENL degradation in both 2D10 and J-Lat A1 cells **(Fig. 1I)**. MS41 is derived from PFI-6, a dual inhibitor targeting the YEATS domains of ENL and AF9 ^18^. Interestingly, although MS41 can degrade ENL, it seems not to be able to degrade AF9, possibly due to the extremely low expression of AF9 in Jurkat cells. Therefore, the observed effects of HIV latency reversal are likely driven primarily by ENL degradation, but not AF9 **(Fig. S1G)**. A similar phenomenon has been observed in leukemia cell lines and several MLL-rearranged leukemia models, where AF9 expression is low or undetectable^18^. Nevertheless, continued efforts are required to develop PROTACs capable of selectively degrading ENL while sparing AF9. With that, we next tested the ENL-specific PROTAC Cpd-1 ^19^, which covalently links thalidomide, a commonly used ligand for the E3 ubiquitin ligase Cereblon. As expected, Cpd-1 significantly enhanced HIV latency reversal by inducing GFP expression in 2D10, J-Lat A1, and J-Lat 10.6 cells in a dose-dependent manner **(Fig. 1J-1L; Fig. S1H-J)** and efficiently induced ENL degradation **(Fig. 1M)**.

Together, these chemical biology approaches demonstrate that suppressing ENL effectively reactivates latent HIV-1 in multiple cellular models, highlighting ENL as a critical regulator of HIV latency and a potential target for latency-reversing strategies. This contrasts with the long-standing hypothesis that ENL acts as an activator within the SEC during HIV transcription.

### Proximity proteomics reveals ENL interactors involved in HIV transcriptional regulation

To identify how ENL regulates HIV latency, we mapped its protein-protein interactome using (Mini) TurboID-based proximity-dependent biotinylation to capture ENL-interacting proteins ^20^. A doxycycline-inducible, full-length ENL fusion construct was stably integrated into Jurkat 2D10 cells **(Fig. 2A)**. Upon doxycycline induction, ENL expression led to decreased GFP levels in 2D10 cells, further supporting that ENL functions as a suppressor rather than an activator of HIV transcription **(Fig. 2B)**. This was accompanied by reduced HIV transcription **(Fig. 2C)** and increased ENL mRNA levels **(Fig. 2D)** without detectable cellular toxicity **(Fig. S1K)**. Proteomics analysis identified 2,037 proteins that significantly interacted with ENL, compared with controls (*p* < 0.05) **(Fig. 2E and Table S1)**, including established ENL interactors such as P-TEFb (CCNT1 and CDK9) and HIV latency–associated factor BRD4 **(Fig. 2F)**. Gene ontology analysis revealed enrichment of proteins involved in histone modification, transcriptional elongation from RNA polymerase II, and regulation of mRNA metabolic process **(Fig. 2G and Table S2)**. Unexpectedly, ENL pulldown revealed significant enrichment of the deubiquitinase USP7, and protein-protein interaction analysis indicated that ENL, BRD4, and USP7 may form a regulatory complex **(Fig. 2H)**. Similarly, ENL interactome proteins were enriched for functional categories related to transcriptional regulation, highlighting the presence of factors implicated in the HIV life cycle **(Fig. 2I)**. These proteins also included bromodomains -containing and RNA-binding domains **(Fig. 2J)**, as well as subunits of the BRD4 and BRD4-P-TEFb complexes **(Fig. 2K)**, representing key components of central transcriptional regulatory hubs.

**Figure 2.**
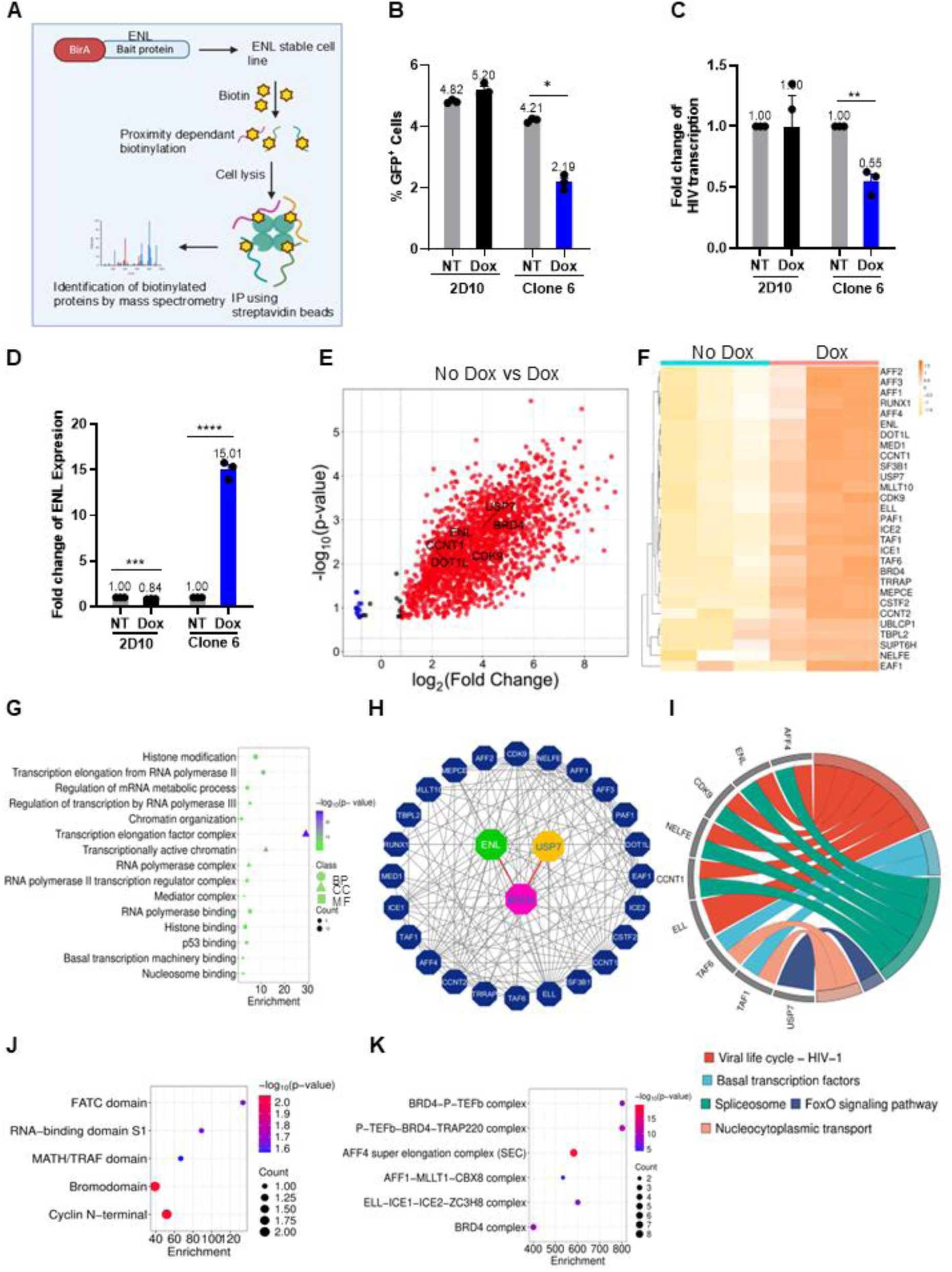
Generation of doxycycline-inducible ENL-expressing stable cell line and quantitative mass spectrometry analysis. Doxycycline-inducible ENL stable cell lines were generated and selected with blasticidin (4 mg/mL) for two weeks. ENL expression was induced with doxycycline (0.4 µg/mL) for 72 h prior to downstream assays. ENL stable cell line harboring the inducible ENL construct or 2D10 cells were induced for three days prior to proximity labeling and affinity purification. A schematic of the proximity biotinylation workflow is shown (A) (created with BioRender). HIV reactivation was quantified by GFP expression using flow cytometry (B), while HIV transcription was measured by qPCR (C). Fold change in ENL expression following doxycycline induction is shown (D). Quantitative mass spectrometry analysis of ENL-stable cells identified SEC-associated proteins, highlighted in black, with enrichment values plotted along the x-axis comparing No Dox to Dox conditions (E). Expression patterns of SEC-associated proteins were visualized as a heatmap using SRplot (F), and functional enrichment was performed using ShinyGO 0.85.1 to assess Gene Ontology categories (G), KEGG pathways (I), InterPro domains (J), and CORUM complexes (K), visualized using SRplot. Protein–protein interaction networks of SEC-associated proteins were generated using Cytoscape (H). Statistical significance for enrichment analyses was determined using a hypergeometric test. Detailed information is given in Table S1.

### Validation of USP7 and ENL in maintaining HIV-1 Latency by CRISPR/Cas9 KO

Our proteomic data support a possible role of USP7 in regulating HIV latency through interaction with the ENL–BRD4–P-TEFb complex, where it may stabilize ENL or other components via its deubiquitinating activity, thereby modulating transcriptional regulation. To test this hypothesis, the USP7-targeting PROTAC CST967 ^21^, a potent and selective CRBN-recruiting degrader, was added to HIV latency models. We found that degradation of USP7 by CST967 efficiently enhanced HIV-driven GFP expression in 2D10 and J-Lat A1 cells, across concentrations ranging from 5 to 20 µM **(Fig. 3A, B; Fig. S1L, M)**, supporting a role for USP7 in suppressing proviral transcription. Maximal USP7 degradation was observed at 24 h across a dose range of 15-20 µM in both models, in a dose-dependent manner **(Fig. 3C)**. To further evaluate the role of ENL or USP7 in HIV transcription, 2D10 cells were electroporated with RNPs carrying sgRNAs targeting ENL or USP7, or a non-targeting control, for CRISPR/Cas9 gene knockout (KO). KO efficiency of ENL/USP7 was quantified by qPCR, confirming a significant reduction of ENL or USP7 relative to the non-targeting control **(Fig. 3D)**, which was then validated by western blotting from nucleofected cells, confirming a near-complete loss of ENL or USP7, without altering Cyclin T1 or CDK9 protein expression **(Fig. 3E)**. CRISPR-mediated knockout of ENL or USP7 in 2D10 cells also led to a significant increase in HIV transcription **(Fig. 3F)**, without detectable cellular toxicity observed **(Fig. 3G)**. Together, these findings support a suppressive role of ENL and USP7 in HIV transcription and further establish the ENL–USP7 axis as a key regulator of HIV latency.

**Figure 3.**
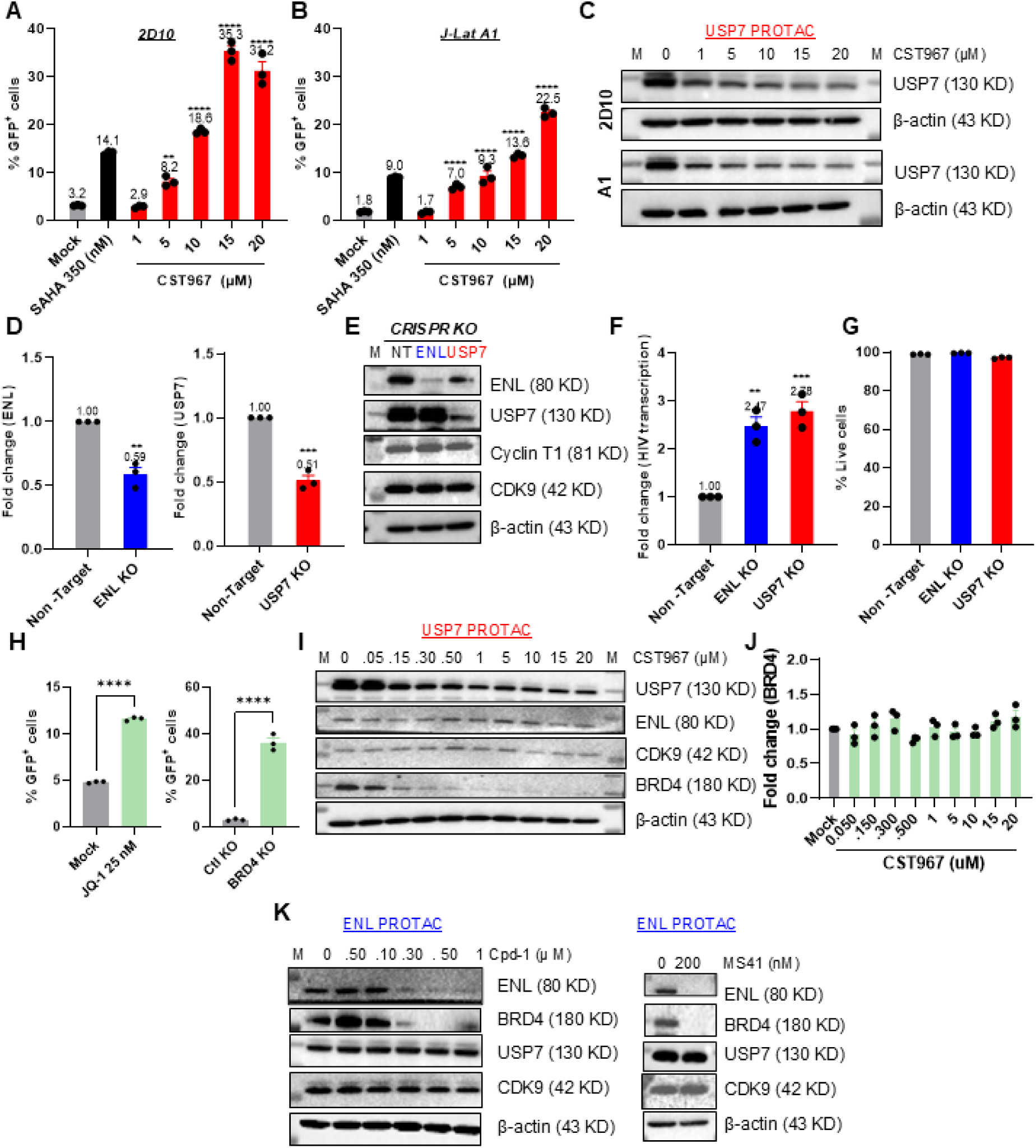
USP7-targeting PROTAC degradation disrupts HIV latency. 2D10 and J-Lat A1 cells were treated with increasing concentrations of the USP7-directed PROTAC CST967. Cells were harvested 24 h post-treatment, and HIV reactivation was quantified by GFP expression using flow cytometry. Dose-dependent GFP induction following CST967 treatment in 2D10 (A) and J-Lat A1 cells (B) is shown; USP7 degradation was verified by immunoblotting (C). CRISPR-mediated knockout of USP7/ENL was performed in 2D10 cells using targeting USP7/ENL or non-targeting control gRNAs, and cells were harvested for five days post-nucleofection. Knockout efficiency was quantified by fold-change in USP7 and ENL transcript levels by qPCR (D). ENL and USP7 depletion was verified by immunoblotting (E) while HIV-1 reactivation was measured by flow cytometry (F). Cell viability was assessed using Live/Dead dye during flow cytometry acquisition (G). 2D10 cells were treated with JQ1 for 24 hours or CRISPR-mediated knockout of BRD4 for 3 days, and HIV-1 reactivation was measured by flow cytometry (H). 2D10 cells were treated with the ENL-targeting PROTACs MS41 and Cpd-1 or with the USP7-targeted PROTAC CST967 for 24 h prior to protein and transcript analysis. Whole-cell lysates were prepared and subjected to immunoblotting to assess protein levels in CST 967 (I) treated cells. BRD4 mRNA abundance in CST967-treated cells was quantified by qPCR (J). Immunoblotting was employed to assess protein levels in Cpd-1 and MS41 treated cells (K). β-Actin served as a loading control. Experiments were performed with three biological replicates per condition. The *P* values were determined using one-way ANOVA or two-tailed Student’s t-test.

### USP7 stabilizes BRD4 through interaction with ENL to control HIV-1 latency

Our chemical biology, genetics, and proteomics data support that ENL forms a complex with USP7, which is critical for HIV latency. However, the underlying mechanism remains unclear. Interestingly, ENL not only pulled down USP7 but also the well-established HIV latency-associated protein BRD4^22–25^. Since USP7 is a protein deubiquitase, we hypothesize that it may protect ENL, BRD4, or both. Consistent with previous reports ^24,26^, either inhibition of BRD4 by its specific inhibitor JQ1 or BRD4 KO significantly induced HIV expression from latency in 2D10 cells **(Fig. 3H)**, confirming that BRD4 suppresses HIV transcription ^25^, and is therefore a bona fide HIV latency factor. It has been shown that BRD4 has several isoforms, due to alternative splicing^27^. To examine which isoforms are present in 2D10 cells, the BRD4 gene was knocked out using CRISPR/Cas9 KO. At least two isoforms were detected (∼ 110-120 KD and ∼180KD), both of which were almost completely lost after KO **(Fig. S2)**. Thus, these data confirm the specificity of the BRD4 antibody used in this study. Treatment of 2D10 cells with USP7 PROTAC CST967 for 24 h effectively depleted USP7 protein while leaving ENL and CDK9 unaffected **(Fig. 3I)**. Of note, the BRD4 protein was almost completely lost by USP7 degradation. Importantly, BRD4 mRNA remained unchanged in USP7 KO 2D10 cells **(Fig. 3J)**. Together, these data strongly support that USP7 acts as a deubiquitinase that stabilizes BRD4. Surprisingly, treatment of cells with ENL PROTACs Cpd-1 and MS41 was also able to effectively reduce BRD4 protein level without affecting USP7 or CDK9 **(Fig. 3K)**. Considering that ENL forms a complex with USP7 and BRD4, along with CDK9 or cyclin T1, i.e., P-TEFb **(Fig. 4A)**, these observations indicate that ENL may recruit USP7 to maintain BRD4 stability.

**Figure 4.**
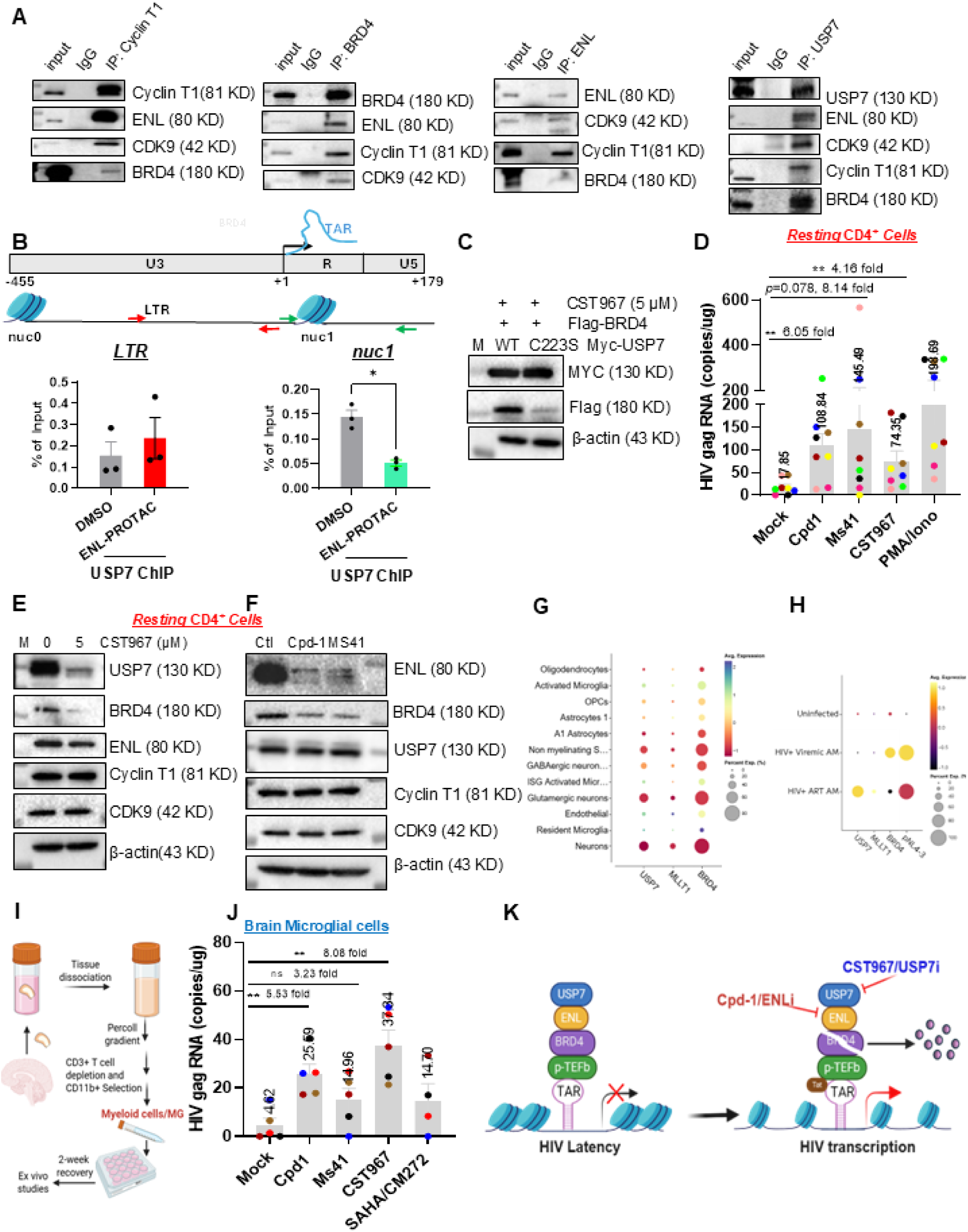
Targeted degradation of USP7 stabilizes BRD4 and promotes HIV latency reversal in resting CD4⁺ T cells and the brain microglia isolated from PWH receiving suppressive ART. Immunoprecipitation was performed using anti-Cyclin T1, anti-BRD4, anti-ENL, or anti-USP7 antibodies, followed by immunoblot analysis to evaluate complex associations in 2D10 cells (A). 2D10 cells were treated with 500 nM Cpd-1 for 24 h. Chromatin immunoprecipitation was performed using a USP7-specific antibody. Enriched DNA fragments were amplified using LTR-and NUC1-specific primers and quantified by SYBR Green qPCR (B). Immunoblot analysis of 293T cells co-transfected with a plasmid expressing Myc-tagged wild-type USP7, or mutant Myc-USP7 C223S, together with Flag-tagged BRD4. Cells were harvested at 48 h p.t and subjected to western blot analysis (C). Resting CD4⁺ T cells were isolated from the peripheral blood of PWH on suppressive ART and treated ex vivo with Cpd-1 (500 nM), MS41 (1,000 nM), CST967 (5 µM), PMA plus ionomycin (200 ng/mL and 2 µM), or DMSO for 24 h. HIV transcriptional induction was quantified by RT-ddPCR, measuring cell-associated HIV gag RNA (D). USP7/ENL degradation was verified by immunoblotting (E and F). Brain tissue samples were obtained from the Last Gift cohort, including altruistic PWH who are viremic or suppressed receiving ART, to profile and integrate CNS cellular transcriptomes. Brain microglia single cells were isolated following mechanical dissociation and enzymatic digestion, and a CNS single-cell suspension was generated after Percoll separation. CD3⁺ T cells were positively selected from the suspension, and dot plot expression of BRD4, MLLT1, and USP7 across the different cell types is shown in CD3⁺ T cells (G). Expression of USP7, ENL, and BRD4 in HIV RNA⁺ activated microglial cells is shown relative to HIV⁻ bystander microglia from uninfected, viremic, and ART-suppressed activated conditions (H). Brain microglia were isolated from the CD3⁻ fraction through CD11b⁺ selection. A schematic layout illustrating the isolation of brain microglia from brain tissues collected from PWH is shown (I). Brain microglia cells were treated ex vivo with Cpd-1 (500 nM), MS41 (1,000 nM), CST967 (5 µM), SAHA (250 nM) plus CM272 (50 nM) or DMSO for 24 h. HIV transcriptional induction was quantified by RT-ddPCR, measuring cell-associated HIV gag RNA (J), and *P* values were determined using one-way ANOVA. Schematic representation of the role of the ENL-BRD4-USP7 complex in regulating HIV-1 latency and transcription is shown (K). Under latent conditions (left), the ENL/USP7/BRD4 complex associates with the HIV promoter and restricts transcriptional elongation, maintaining viral latency. Pharmacological targeting of ENL (Cpd-1) or USP7 (CST967) disrupts this complex, leading to induction of HIV transcription. Mechanistically, USP7 stabilizes BRD4 through its deubiquitylating activity, thereby sustaining latency. Disruption of USP7 function reduces BRD4 stability, promoting transcriptional activation. The right panel illustrates the proposed model in which loss of USP7 activity destabilizes BRD4 and alters ENL-mediated regulation, resulting in enhanced HIV transcription.

To further determine whether ENL mediates USP7 recruitment into the HIV promoter, cells were treated with the ENL-targeting PROTAC Cpd-1, followed by ChIP using a USP7-specific antibody. We found that depletion of ENL significantly reduced USP7 recruitment to the nuc1 region but not the LTR region of the HIV promoter **(Fig. 4B)**. To determine whether BRD4 protein stabilization is dependent on USP7 deubiquitylating activity, 293T cells were treated with CST967 for 24 hours and then co-transfected with plasmids expressing either WT-USP7 or the catalytically inactive mutant USP7-C223S ^28^ with WT-BRD4. Overexpression of WT-USP7, but not the C223S mutant, increased BRD4 protein levels, proving that the stability of BRD4 is dependent on USP7 deubiquitylating activity **(Fig. 4C)**. Collectively, these findings demonstrate that ENL organizes a BRD4–USP7 chromatin regulatory module at the HIV LTR during HIV latency, where USP7 stabilizes BRD4 to maintain proviral quiescence.

### Suppressing the ENL-USP7 axis reactivates latent HIV in resting CD4⁺ T cells

Targeting ENL or USP7 enhanced HIV reactivation in latency models *in vitro*, prompting us to validate their roles in resting CD4⁺ T cells from PWH *ex vivo*. Peripheral blood was obtained from eight antiretroviral therapy (ART)-suppressed PWH with undetectable viral loads (**Table S3**). Resting CD4⁺ T cells were then isolated and treated for 24 h with DMSO, PMA/ionomycin (positive control), Cpd-1, MS41, or CST967. HIV transcription following reactivation was quantified by digital droplet PCR (ddPCR) targeting the gag region of the viral genome. Treatment with Cpd-1 or MS41 significantly induced HIV transcription in CD4⁺ T cells from 7 of 8 participants, while CST967 activated HIV transcription in 6 of 8 patients, with robust reactivation observed across compounds: Cpd-1 6.05-fold, MS41 8.14-fold, and CST967 4.16-fold **(Fig. 4D; Fig. S3)**. As observed in Jurkat models, CST967 and Cpd-1/MS41 efficiently degraded USP7 **(Fig. 4E)** and ENL **(Fig. 4F)**, respectively, without affecting Cyclin T1 or CDK9 in resting CD4⁺ T cells. Notably, BRD4 was nearly abolished in both conditions, supporting a model in which ENL recruits USP7 to stabilize BRD4. Interestingly, there was a modest reduction of ENL protein by CST967, indicating that USP7 may also contribute to ENL protein stability. These findings indicate that perturbing ENL or USP7 effectively relieves transcriptional blocks imposed during latency, thereby reactivating latent HIV in resting CD4⁺ T cells, further confirming their essential roles in HIV latency.

### Single-nuclei RNA-seq reveals persistent regulator expression sustaining HIV latency in the brain microglia

In addition to the T cell reservoir, we recently characterized a stable HIV reservoir in human brain myeloid cells/microglia (MG)^29^. We next asked whether ENL or USP7 plays a similar role in myeloid HIV reservoirs. Thus, we performed single-nuclei RNA sequencing (snRNA-seq) on brain tissues to profile and integrate CNS cell transcriptomes from individuals without HIV, viremic PWH, and PWH receiving suppressive ART. Clustering of single-nucleus transcriptomes resolved the CNS into distinct populations, including microglia, astrocytes, and oligodendrocytes **(Fig. 4G)**. Notably, key HIV latency regulators such as BRD4, ENL (MLLT1), and USP7 exhibited pronounced heterogeneity across CNS cell clusters **(Fig. S4A)**, with increased expression following ART **(Fig. S4B).** Expression of these regulators was distributed across cell types, with the highest levels observed in the activated microglia subset. Within the activated microglial subset, the expression of BRD4, ENL, and USP7 was elevated during viremic infection; however, levels did not return to baseline following ART, indicating that HIV latent infection is associated with persistent upregulation of these HIV latency factors even under viral suppression **(Fig. S4C)**. In ART-suppressed activated microglia, HIV RNA⁺ cells display elevated USP7 and MLLT1 expression relative to HIV- bystander cells, and to cells derived from viremic and uninfected conditions **(Fig. 4H)**. Further, the number of ENL⁺ **(Fig. S4D)**, BRD4⁺ **(Fig. S4E)**, and USP7⁺ **(Fig. S4F)** microglia and astrocytes was higher in ART-suppressed samples compared with viremic and HIV-negative controls, highlighting a possible role of brain microglia, and presumably astrocytes, in shaping HIV latency regulators, i.e., ENL, BRD4, and USP7, during latent HIV infection in CNS-resident immune cells.

To investigate the contribution of the ENL/USP7/BRD4 axis to HIV latency in the brain myeloid cells/MG, we cultured human myeloid cells/MG isolated from ART-suppressed individuals **(Fig. 4I and Table S4)** and treated them with the above-characterized ENL or USP7 inhibitor or PROTACTs for 24 hours. We found that the treatment of ART-suppressed human MG with these small molecules elicited robust HIV reactivation, where the ENL PROTAC Cpd-1 induced a 5.53-fold increase across all five donors, MS41 triggered a 3.23-fold increase in 4/5 donors, and the USP7 PROTAC CST967 produced an 8.08-fold induction across all donors **(Fig. 4J; Fig. S5A)**. Consistent with findings in resting CD4⁺ T cells, we observed that CST967 induced efficient degradation of USP7 in brain MG **(Fig. S5B)**, indicating that, similar to T cells, ENL/USP7 likely participates in HIV latency in the brain myeloid cells/MG. Taken together, these data reveal that the key ENL-USP7 axis at the SEC sustains HIV latency in the brain myeloid cells/MG, in addition to resting CD4⁺ T cells.

## Discussion

Efforts to cure HIV-1 are hindered by the ability of replication-competent proviruses to switch between latent and transcriptionally active states after treatment interruption, a process governed by inducible transcriptional regulators. HIV latency is reinforced in part by blocks to transcriptional elongation, which has motivated latency-reversal strategies aimed at releasing P-TEFb from inhibitory complexes such as 7SK snRNP and BRD4 ^22,30^. A substantial fraction of cellular P-TEFb is also sequestered within the SEC, which coordinates inducible gene expression^27^. Although ENL is well recognized as a crotonylation/acetylation reader, prevailing models further propose that it promotes SEC assembly and active transcriptional elongation. However, whether this activity is mechanistically linked to its function as an epigenetic reader remains unclear.

Notably, some transcriptional regulators exert opposite effects on host versus HIV gene expression; BRD4 is one such example. Here, using complementary chemical biology, proteomic, and genetic approaches, we identify ENL as another factor with this paradoxical behavior. In contrast to its proposed role as a transcriptional activator, ENL functions similarly to BRD4 as a suppressor rather than an activator of HIV transcription across multiple in vitro and ex vivo latency models, including brain myeloid cells and microglia. Although ENL was previously proposed to activate HIV transcription as part of the SEC at the HIV promoter ^31^, strong experimental support for this model has been limited. Instead, emerging studies suggest that the SEC can negatively regulate HIV transcription. Our findings provide a molecular explanation for this unexpected activity by showing that the ENL–USP7 regulatory axis controls BRD4 protein stability to enforce HIV latency. Together, these data suggest that ENL may be a key determinant of the SEC’s latency-promoting function.

Previous studies have shown that histone crotonylation regulates HIV latency through the fatty acid-metabolizing enzyme ACSS2 ^32^, enhances IAPi (AZD5582)-mediated HIV latency reversal and promotes the cleavage of p100 into p52 ^33^, collectively highlighting histone lysine crotonylation as an epigenetic regulator of HIV transcription with potential relevance to eradication strategies. These observations prompted us to test whether ENL regulates HIV latency through its canonical acylation-reader function, potentially by recognizing crotonylated proteins within the SEC. Proximity-dependent biotinylation proteomics identified several candidate crotonylated proteins within the ENL interactome. However, CRISPR knockout of selected candidates, including UBN1, ZMYND8, and ASXL3, did not reveal a role in HIV transcription (data not shown). Together, these results suggest either that ENL recruitment into the SEC does not depend on its epigenetic reader function in this context or that the relevant acylation-dependent mechanism remains to be identified.

Our study also identifies USP7 as a critical deubiquitinase for BRD4. However, the E3 ubiquitin ligase that counterbalances USP7 to control BRD4 stability within this regulatory network remains unknown. SPOP (speckle-type POZ protein), a known E3 ubiquitin ligase adaptor, has been reported to regulate BRD4 turnover in epithelial cells ^34^, but SPOP was not detected in our T cell proteomic dataset. Although SPOP has been implicated in ubiquitin-dependent BRD4 degradation, its effects may be context-dependent and, in some settings, associated with BRD4 stabilization ^34^. We also identified several ubiquitin-related proteins in the ENL pulldown dataset, including β-TrCP and UBA (**Fig. S6**), but immunoblot analyses did not support a direct role for these factors in BRD4 degradation in T cells. Defining the ligase machinery that opposes USP7 will therefore require further investigation.

Unlike the broad-spectrum DUB inhibitor PR-619, which nonspecifically targets multiple enzymes, including USP7 and USP47 ^35^, CST967 is a selective USP7 degrader that suppresses HIV transcription. Selective PROTAC-mediated degradation of USP7 or ENL caused a marked loss of BRD4 without affecting Cyclin T1 or CDK9, supporting the existence of a selective regulatory axis in Jurkat latency models *in vitro* and in resting CD4⁺ T cells *ex vivo*. These findings support a model in which ENL recruits USP7 to preserve BRD4 stability and maintain transcriptional incompetence in resting CD4⁺ T cells. Consistent with this interpretation, only wild-type USP7, but not the catalytically inactive C223S mutant, increased BRD4 protein abundance, indicating that USP7-dependent stabilization of BRD4 requires its deubiquitinase activity. More broadly, USP7 has been shown to stabilize multiple transcriptional regulators through ubiquitin-dependent mechanisms, including NOTCH1 ^36^, ZNF638 ^37^, and Nrf1a ^38^. Our results extend this paradigm to BRD4 in the setting of HIV latency.

In the current study, single-nucleus RNA-seq further suggests that persistent expression of the ENL–USP7–BRD4 axis may contribute to the maintenance of HIV latency in microglia, consistent with the significant latency reversal observed when ENL or USP7 was inhibited in microglia isolated from ART-suppressed people with HIV. Our previous work likewise showed that brain microglia persist as an HIV reservoir during suppressive ART ^29^. Nevertheless, the detailed molecular basis of this pathway in microglia remains incompletely defined. Progress in this area is complicated by the lack of a physiologically relevant microglial latency model suitable for mechanistic study which represents an important limitation that future work will need to address. Together, our findings demonstrate that ENL acts as a suppressor rather than an activator of HIV transcription and that ENL cooperates with USP7 to stabilize BRD4 and maintain this latency-promoting axis **(Fig. 4K)**. Targeting ENL or USP7 with small molecules or PROTACs disrupts this pathway, destabilizes BRD4, and reactivates latent HIV in both T cells and brain myeloid cells, including microglia. These findings uncover an unexpected role for the ENL–USP7–BRD4 axis and for SEC-associated transcriptional suppression in sustaining HIV latency, thereby identifying this pathway as a potentially actionable target for HIV eradication strategies.

## Methods

### Cell culture and HIV-1 latency models

Jurkat-derived HIV-1 latency models (2D10, J-Lat A1, J-Lat 10.6) were maintained in RPMI 1640 medium supplemented with 10% fetal bovine serum and sodium pyruvate, 2 mg/mL L-glutamine, 10mM HEPES and penicillin-streptomycin at 37°C with 5% CO₂ as described earlier ^33,39^. J-Lat A1 and J-Lat 10.6 cells were obtained from the NIH AIDS Reagent Program, and 2D10 cells were generous gift by Jonathan Karn (Case Western Reserve University). 293T cells were maintained in DMEM (Thermo Fisher Scientific, USA) supplemented with 10% FBS.

### Latency reversal agents

For HIV-1 latency reversal, cells were treated with the ENL PROTAC MS41 ^18^, Cpd-1 ^19^, ENL inhibitors ENLi13 and SR0813 ^16,17^, and USP7 PROTAC (CST967) ^40^, as detailed in the Reagents Table S5.

### Quantification of HIV-1 expression by qPCR

Total RNA was extracted using the RNeasy Mini Kit (Qiagen) and treated with DNase I (Invitrogen) to remove genomic DNA. First-strand cDNA was synthesized from the purified RNA using SuperScript III reverse transcriptase (Invitrogen) with random primers (Invitrogen). TaqMan real-time PCR was performed on an ABI QuantStudio 5 system, with HIV-1 transcripts amplified using primers and probes specific for the gag region of HIV as described earlier ^29,41^. The SDHA probe (Thermo Scientific) was used as a control for real-time PCR, where indicated.

### Quantification of Cell-Associated HIV RNA by ddPCR

Cell-associated HIV RNA was measured by ddPCR. Total RNA was isolated from PWH-derived resting CD4⁺ T cells and brain microglia using the RNeasy Mini Kit (Qiagen), treated with DNase I, and reverse transcribed with SuperScript IV (Invitrogen). HIV transcripts were quantified by RT-ddPCR using two primer/probe sets targeting HIV *gag* (Table S4). Cycling conditions were 95°C for 10 min, 45 cycles of 94°C for 30 s and 57°C for 60 s, followed by 98°C for 10 min. Droplets were read and analyzed in QuantaSoft using absolute quantification.

### Flow Cytometry

2D10, J-Lat A1, and J-Lat 10.6 cells (5× 10^5^) were treated with PROTACs or inhibitors for 24 h. Cells were harvested and washed with PBS and resuspended in FACS buffer (PBS with 2% FBS). GFP-expressing HIV-1 was quantified on a BD LSRFortessa cytometer (BD Biosciences), with at least 10,000 events recorded per sample. Cell viability was assessed using Live/Dead dye (Invitrogen). Data were analyzed using FlowJo v10 (BD Biosciences).

### CRISPR/Cas9-mediated knockout of host genes in HIV latency models

Pre-designed CRISPR RNAs (crRNAs) targeting ENL and USP7, along with scrambled non-targeting controls (NT), were purchased from Integrated DNA Technologies (IDT). Sequences of all crRNAs used are listed in the Reagent Table S4. CRISPR/Cas9 ribonucleoprotein (RNP) complexes were prepared, crRNA/tracrRNA duplexes were annealed, and electroporation was performed as described previously ^42^. Three crRNAs targeting different regions of the gene were multiplexed to improve knockout efficiency. For each electroporation, 2 × 10⁶ latently infected cell lines were washed twice with PBS (90 × g, 10 min) and resuspended in SE Cell Line Nucleofector Solution (Lonza) at 1 × 10⁶ cells per 20 μL. Resuspended cells were combined with crRNP complexes and immediately transferred into cuvettes of the P3 Primary Cell Nucleofector Kit (Lonza) and electroporated using code CM-120 on the Lonza 4D-Nucleofector. Following electroporation, cells were resuspended in 200 μL of prewarmed, supplemented RPMI and cultured at 37 °C. Cells were maintained at 1 × 10⁶ cells/mL and provided with fresh media as needed before downstream experiments.

### Plasmids Transfection

293T cells were transfected with plasmid expressing Myc-tagged wild-type USP7 (USP7-WT), mutant Myc-USP7 C223S (USP7-C223S), or Flag-tagged BRD4 (BRD4-WT) (Addgene; Table S4). Cells were seeded in six-well plates and, the following day, treated with CST967 PROTAC (5 μM). After 24 h of treatment, cells were transfected using Lipofectamine 3000 transfection reagent (Invitrogen, USA) according to the manufacturer’s instructions. Briefly, 2 μg of each plasmid DNA was mixed with the transfection reagent following their recommended protocol, and the complexes were added to the cells. At 48 h post-transfection, cells were washed once with PBS, harvested, and subjected to western blot analysis.

### Immunoblot analysis

HIV latency model cells (1 × 10⁶) were treated with PROTACs for 24 h. For CRISPR/Cas9-mediated knockout experiments, cells were harvested up to 5 days post-nucleofection. Whole-cell and nuclear proteins were extracted in RIPA buffer (Sigma) containing protease and phosphatase inhibitors (Cell Signaling). Protein concentrations were determined using the BCA Protein Assay Kit (Thermo Fisher) according to the manufacturer’s instructions. Equal amounts of protein were resolved by SDS-PAGE (4-12% Tris-glycine gels), transferred to PVDF membranes, and incubated with primary antibodies overnight at 4 °C. After three washes with 1% TBST, membranes were incubated with secondary antibodies for 1 h at room temperature, washed three times, and visualized using chemiluminescence (Thermo Fisher). Protein expression was assessed using the following primary antibodies: anti-ENL, anti-Cyclin T1, anti-CDK9, and anti-β-actin (all from Cell Signaling), anti-BRD4 (Cell signaling for immunoprecipitation or Abcam for western blot) and anti-USP7 (Bethyl).

### Co-immunoprecipitation

Immunoprecipitations (IP) were performed as previously described ^33^. Briefly, cells were harvested, washed with PBS, and Nuclear and cytoplasmic proteins were separated with NE-PER™ reagents (Thermo Scientific) per manufacturer’s instructions. Cells were sonicated briefly (5 pulses, repeated three times) and centrifuged at 10,000 × g for 10 min at 4 °C. Supernatants were collected, protein concentrations measured by BCA, and normalized to equal amounts. Samples were incubated with the appropriate antibody (dilutions as recommended by the manufacturer) on a rocker overnight at 4 °C. Protein-antibody complexes were captured with 40 µL Protein A/G beads (Thermo Scientific), washed three times with lysis buffer, and eluted for downstream analysis.

### Chromatin immunoprecipitation (ChIP) assays

ChIP assays were carried out as described earlier ^43^ with modifications. 2D10 cells were treated for 24 hours with ENL-targeting PROTAC (Cpd-1 500 nM) or with DMSO. Five million cells were cross-linked with 1% formaldehyde for 10 min at room temperature, then quenched with 125 mM glycine. Cells were washed with PBS, flash-frozen in liquid nitrogen, and stored at -80°C. Nuclei were isolated by resuspending the pellet in 10 mL rinse buffer #1, incubating on ice for 10 min, and centrifuging at 1,200 × g, 5 min, 4°C followed by resuspension in 10 mL rinse buffer #2 and a second centrifugation. Nuclei were washed twice with 5 mL shearing buffer (1,200 × g, 3 min, 4°C) to remove residual salts, then resuspended in 90 μL shearing buffer with protease inhibitors and transferred to Covaris glass microtubes. Chromatin fragmentation was performed by adding 10 μL of MegaShear nanodroplet cavitation reagent (Triangle Biotechnology) to each tube, followed by acoustic shearing in a Covaris E110 instrument at 4 °C for 7.5 min. The fragmented chromatin was subsequently subjected to immunoprecipitation using antibodies specific for USP7 (Bethyl), with normal rabbit IgG (Cell Signaling) used as a control. DNA associated with the immunoprecipitated complexes was purified using the ChIP Clean & Concentrator kit (Zymo Research). The recovered DNA was quantified by SYBR Green-based qPCR (Applied Biosystems) using primers specific for the HIV LTR and Nuc1 regions. Enrichment was calculated relative to input DNA.

### Isolation and culturing of resting CD4⁺ T cells and brain microglia for HIV latency reversal studies

Resting CD4⁺ T cells were isolated from the peripheral blood of PWH (22-62 years) on suppressive ART using the EasySep kit (STEMCELL Technologies) as previously described ^29,32^. Isolation and culture of brain microglia were performed following a previously established protocol ^29,44^. Briefly, brain tissues from individuals on ART were obtained within 6 h of death from donors enrolled in the “Last Gift” cohort. Within 24 h, tissue samples from the indicated brain regions were mechanically and enzymatically dissociated to generate a single-cell suspension via Percoll separation. CD3⁺ T cells were positively selected for CNS T cell isolation, and BrMCs then were isolated from the CD3⁻ fraction using CD11b⁺ selection. Purified brain microglia (1 × 10⁵) were plated and treated for 24 h with DMSO, 200 ng/mL PMA plus 2 µM ionomycin, SAHA/CM272, or PROTACs targeting ENL (MS41 or Cpd-1) or USP7 (CST967). Cell pellets were collected for RNA isolation, and HIV transcription was quantified using digital droplet PCR (ddPCR) as previously described ^29^.

### Generation of stable cell lines

ENL-stable cell lines with doxycycline-inducible ENL expression were generated as described previously ^20^. Briefly, 2D10 cells were nucleofected with a transposase expression vector (VectorBuilder; pXLone-HA/Tet3G-3xFLAG/miniTurbo:Linker:hMLLT1) together with a piggyBac plasmid. Following nucleofection, cells were selected with blasticidin (4 mg/mL) for two weeks. ENL expression was induced with doxycycline (0.4 µg/mL) for 72 h prior to downstream analyses.

### Proximity-dependent biotinylation and affinity purification

Cells harboring a doxycycline-inducible ENL construct were induced for three days prior to proximity labeling and affinity purification ^20,45^. Corresponding cells maintained without doxycycline were included as controls. Proximity labeling and streptavidin-based affinity purification were executed as described ^20^. Briefly, cells were incubated with 50 µM biotin for 30 min at 37 °C. Biotinylation was induced by exposure to 1 mM H₂O₂ and immediately quenched by washing the cells with a quenching buffer. Cells were resuspended in lysis buffer comprising 50 mM Tris-HCl (pH 7.5), 200 mM NaCl, 10% glycerol, 0.25% Triton X-100, 1 mM MgCl₂, 1 mM PMSF, and supplemented with protease and phosphatase inhibitors. Lysates were rotated for 2 h, then briefly sonicated and centrifuged at 13,000 × g for 20 min. Protein lysates were dialyzed overnight against lysis buffer with two buffer changes to remove excess free biotin. The lysates were clarified by centrifugation at 13,000 × g for 20 min and subsequently incubated with equilibrated streptavidin magnetic beads (Pierce) for 4 h. Beads were rinsed four times with lysis buffer and subsequently four times with 50 mM ammonium bicarbonate (ABC) buffer (pH 7.8). The final bead pellet was resuspended in 50 µL ABC buffer (pH 7.8). The eluates were resolved on a 4-15% gradient SDS-PAGE gel for approximately 1 cm, or until the 25 kDa band became visible. The SDS-PAGE gel was stained with Bio-Safe Coomassie Premixed Staining Solution (BioRad), relevant lanes were excised, and finally processed at the UNC Michael Hooker Proteomics Center, where on-bead trypsin digestion and LC-MS/MS analysis were performed.

### LC-MS/MS and Proteomic data analysis

Following injection onto an Easy Spray PepMap C18 column (75μm id 588 × 25 cm, 2μm particle size), samples were analyzed by LC-MS/MS on an Easy nLC 1200 coupled to Qexactive 587 HF (Thermo Scientific). Mass spectrometry raw data were analyzed with MaxQuant (v1.6.15.0) for protein identification and label-free quantification (LFQ). Peptide-spectrum matches were searched against the reviewed human UniProt database (UP000005640), including an appended common contaminants database (∼250 sequences), using the Andromeda search engine integrated in MaxQuant. Searches were performed using a lysine crotonylation, a fixed modification, while methionine oxidation, cysteine carbamidomethylation and protein N-terminal acetylation were included as variable modifications. Peptide identification employed a target-decoy approach, with false discovery rates set at 1% for both peptides and proteins. Peptides of at least seven amino acids were considered, with up to two missed cleavages permitted. Precursor ions were allowed an initial mass deviation of 7 ppm, and fragment ions a maximum deviation of 0.5 Da. Prior to downstream analysis, proteins representing common contaminants, reverse hits, single-peptide identifications, or with ≥50% missing values were excluded. The averaged log₂ LFQ intensities were computed relative to the corresponding controls, comparing ‘No doxycycline’ and ‘Doxycycline’ conditions. Protein abundances were quantified across biological replicates, and two-sample t-tests were used to evaluate the significance of observed fold-changes. Data visualization and plotting were performed with the SRplot, while protein-protein interaction (PPI) networks were constructed and analyzed in Cytoscape.

### Single-Nuclei Data Integration of CNS Cells from People with HIV on ART

For library preparation and sequencing, flesh frozen, dissected brain tissues were dissociated using Dounce homogenization in nuclei lysis buffer (Sigma, NUC-201) into nuclei, followed by Percoll gradient purification. After dissociation, the filtered homogenates were overlaid by 1.8 M sucrose and centrifuged at 13,000 g for 45 min at 4 °C. The nuclear pellet was aggregated and resuspended in nuclear buffer with RNase inhibitor. Single-nuclei suspensions were filtered through 40 µm strainers and used for 10x Genomics 3′ library preparation.

#### Mapping and counting of snRNA-Seq reads

Read mapping and counting of raw FASTQ data were performed using Cell Ranger v4.0 (10x Genomics) against a combined GRCh38 and HIV reference genome with default parameters, including MAPQ adjustment, transcriptome compatibility, UMI counting, and cell calling. The data is available in BioProject ID PRJNA1357854. The filtered feature-barcode matrix was used for downstream analyses.

Cell clustering and annotation: snRNA-seq data were processed with Parse Biosciences Trailmaker. Low-quality cells (<200 genes, >3% mitochondrial transcripts, or doublets) were excluded. PCA on 2,000 highly variable genes was performed in Seurat, excluding ribosomal, mitochondrial, and cell-cycle genes. Cells were clustered by Louvain and visualized in UMAP (min. distance 0.3, cosine metric).

#### Classification of cell populations

The human brain comprises diverse cell types with distinct molecular and functional profiles. Single-nuclei sequencing enables high-resolution characterization by measuring thousands of transcripts per nuclei. Here, we used a human brain-specific sn-RNA seq reference dataset to annotate clusters and identify representative markers using the MaqQuery function in Seurat v4.3.0.

Differential gene expression: Differentially expressed (DE) genes from each cluster (log₂ fold change ≤ -0.5 or ≥ 0.5, p < 0.05) were further analyzed.

### Statistics

Data are shown as mean ± SEM of at least three independent experiments. Statistical significance was determined by two-tailed Student’s t-test or one-way ANOVA; p < 0.05 was considered significant (*).

### Data Availability

All data are available within this manuscript or its supporting files. Sequencing data generated in this study is available in BioProject ID PRJNA1357854.

## ACKNOWLEDGMENTS

We thank Dr. Brian D. Strahl for helping with the Proximity-dependent biotinylation pulldown assay. We thank Dr. Edward Browne for helping with CRISPR- KO studies. G.J. is supported by NIAID (R21 AI167709-01A1 and R01AI186609), NIMH (R21MH128034 and R01MH136852), CARE (UM1AI164567), and the Collaborative Development Program at B-HIVE (U54AI170855). W.R.L. is supported by NIH grants R35GM145351 and R01CA291968. S.G. is supported by Translational Virology Core at the San Diego Center for AIDS Research (P30AI036214) and The James B. Pendleton Charitable Trust, the “Last Gift” cohort (P01AI169609). J.J. acknowledges the support of the grant R01CA260666 from the National Institutes of Health (NIH) and an endowed professorship by the Icahn School of Medicine at Mount Sinai. This work utilized the NMR Spectrometer Systems at Mount Sinai acquired with funding from NIH SIG Grants 1S10OD025132 and 1S10OD028504. The authors alone are responsible for the content, which does not necessarily reflect the official views of the National Institutes of Health.

## AUTHOR CONTRIBUTIONS

GJ designed the study. W.A performed most of the experiments in this study. XL helped with ENL Inhibitor studies in 2D10 cells and established ENL stable cell lines. SG coordinated rapid research autopsies of human brain tissues. CSS helped with the analysis of single-cell nuclei data, and HC helped perform immunoprecipitation. NK and YT helped isolate brain microglia for snRNA-seq and HIV latency reversal studies. WRL provided the ENL inhibitor (ENLi-13), XL, YS provided ENL PROTAC Cpd-1. KL, HUK, and JJ provided the ENL PROTAC MS41. JK, NMA processed and provided resting CD4+ T cells from ART-suppressed PWH. DMM provided necessary support for this study. WA and GJ prepared the initial draft. GJ processed and finalized the data, compiled the manuscript, and performed multiple revisions. All authors reviewed and approved the final manuscript for submission.

## Declaration of Interests

JJ is a cofounder and equity shareholder in Cullgen, Inc. and Valenyx Therapeutics, Inc., was a scientific cofounder and scientific advisory board member of Onsero Therapeutics, Inc., and is/was a consultant for Cullgen, Inc., EpiCypher, Inc., and Accent Therapeutics, Inc. The Jin laboratory received research funds from Celgene Corporation, Levo Therapeutics, Inc., Cullgen, Inc., and Cullinan Therapeutics, Inc.

---

**Figure S1.**
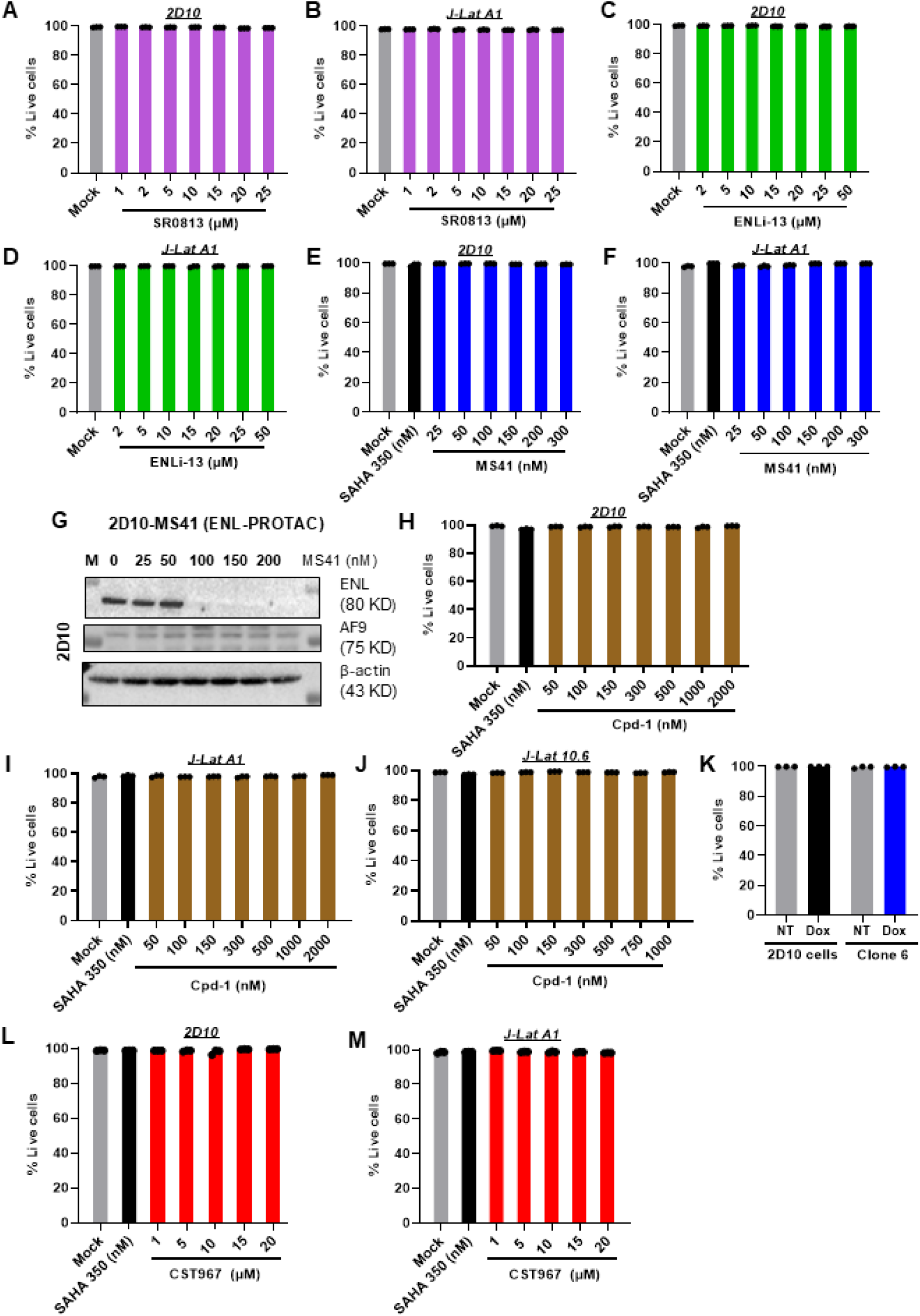
Cell viability following ENL inhibition or degradation in latently infected cell models. Jurkat latency models (2D10, J-Lat A1, and J-Lat 10.6) were treated with indicated concentrations of the ENL inhibitors SR0813 or ENLi-13, or the ENL-targeting PROTACs MS41 and Cpd-1 for 24 h. Cell viability was assessed by flow cytometry using Live/Dead staining. Viability of 2D10 and J-Lat A1 cells following treatment with SR0813 (A and B), ENLi-13 (C and D), and MS41 (E and F), while ENL/AF9 degradation was verified by immunoblotting (G). Cell viability of 2D10, J-Lat A1, and J-Lat 10.6 cells following treatment with Cpd-1 is shown (H-J). Doxycycline-inducible ENL stable cell lines were generated and selected with blasticidin (4 mg/mL) for two weeks, and ENL expression was induced with doxycycline (0.4 µg/mL) for 72 h prior to analysis. Cell viability was assessed by flow cytometry using Live/Dead staining (K). 2D10 and J-Lat A1 cells were treated with increasing concentrations of the USP7-targeting PROTAC CST967 for 24 h, and cell viability was measured in 2D10 (L) and J-Lat A1 cells (M).

**Figure S2.**
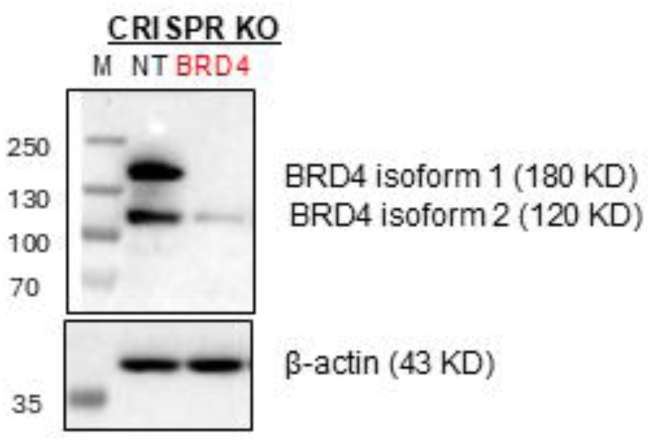
CRISPR-mediated knockout validates the specificity of anti-BRD4 antibodies. CRISPR-mediated knockout of BRD4 was performed in 2D10 cells using targeting BRD4 or non-targeting control gRNAs, and cells were harvested for five days post-nucleofection. Near complete loss of BRD4 validated the specificity of the anti-BRD4 antibody (Abcam) used in this study.

**Figure S3.**
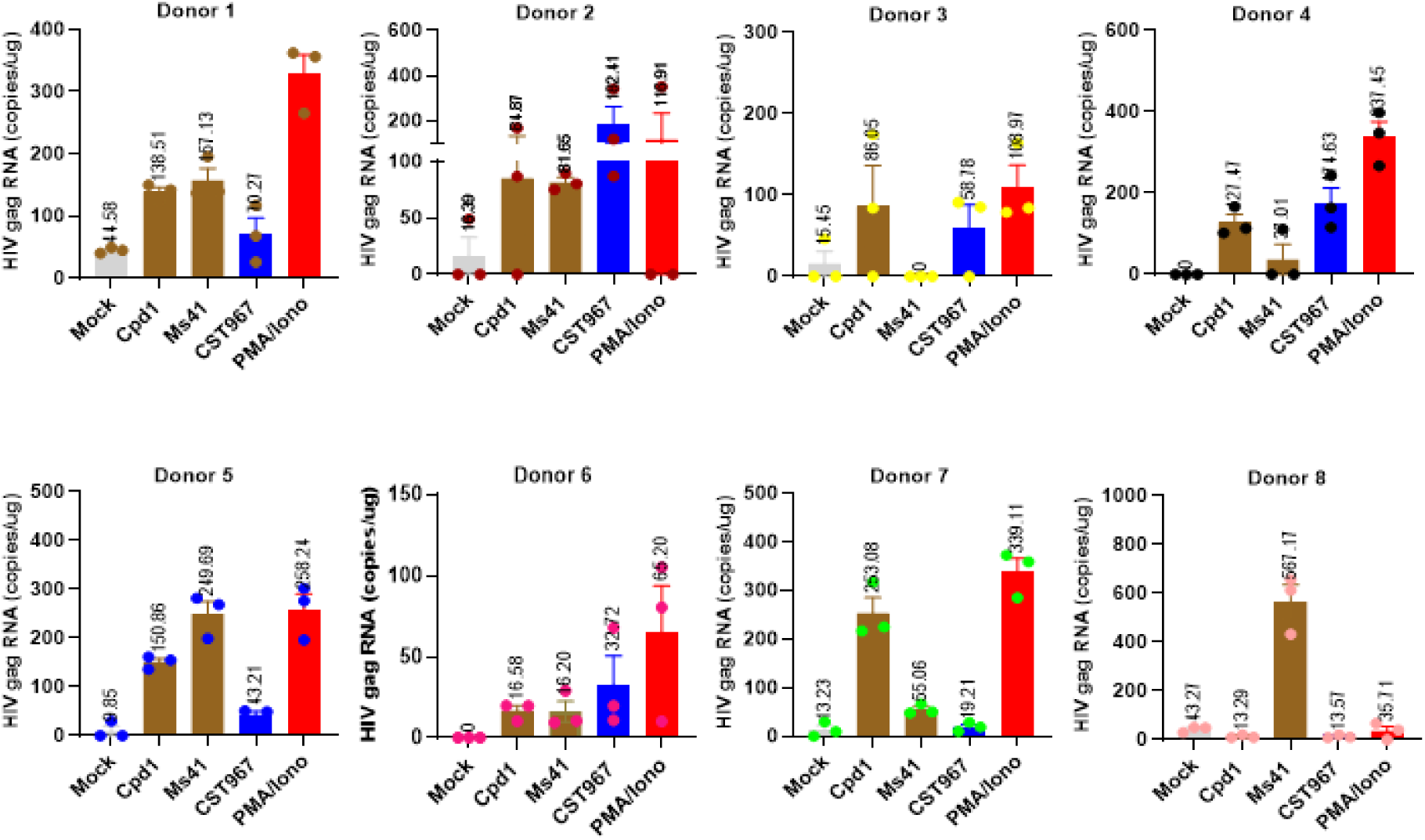
Targeting ENL or USP7 triggers HIV latency reversal in resting CD4⁺ T cells from PWH receiving suppressive ART. Resting CD4⁺ T cells were isolated from the peripheral blood of 8 people with HIV (PWH) on suppressive antiretroviral therapy and treated with Cpd-1 (500 or 1,000 nM), MS41 (1,000 nM), CST967 (5 µM), PMA plus ionomycin (200 ng/mL and 2 µM), or DMSO for 24 h. Data represent individual donor responses. Cell-associated HIV *gag* levels were quantified by ddPCR, and results are presented for each donor with triplicate replicates.

**Figure S4.**
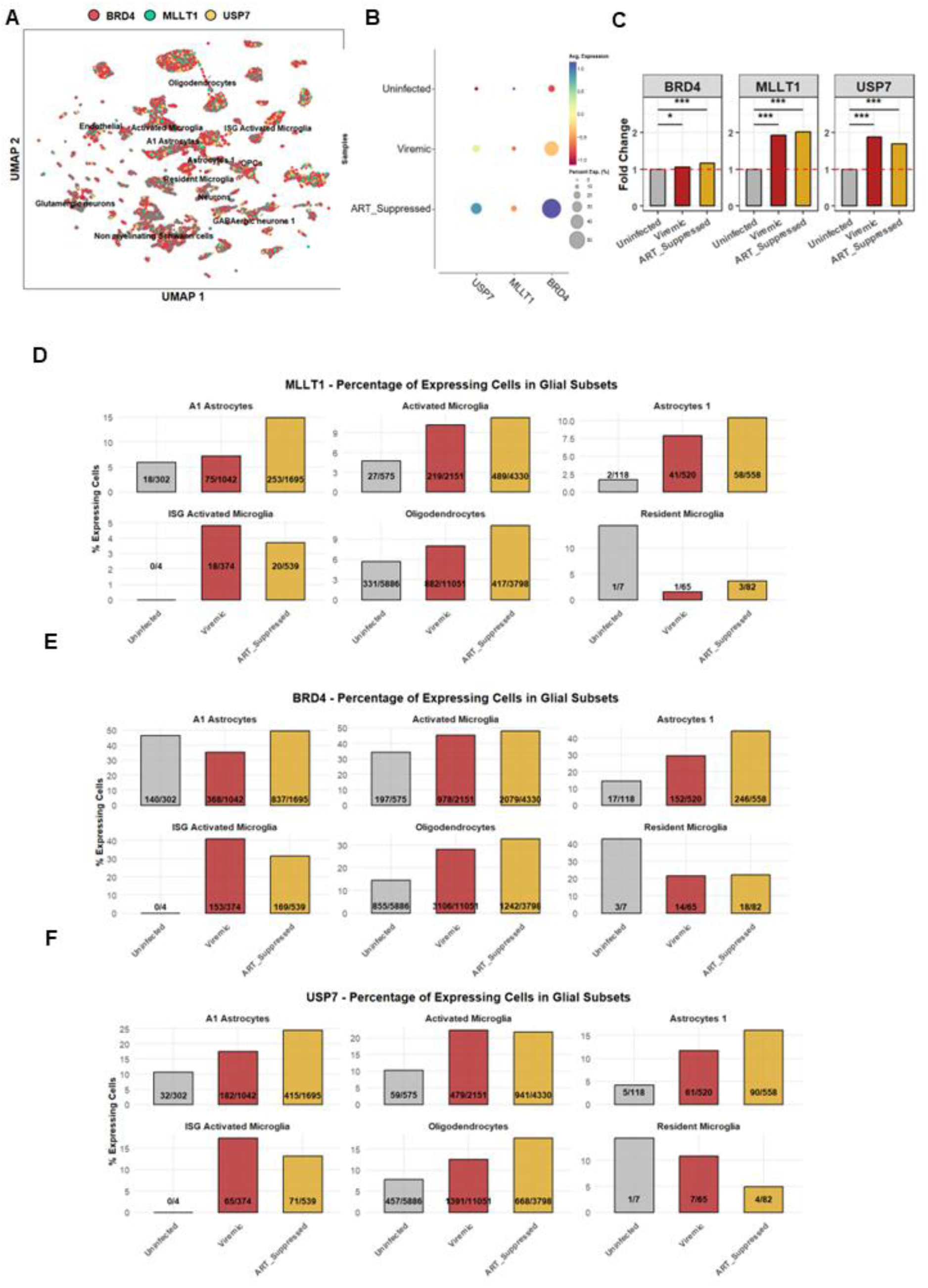
SnRNA-seq analysis links the ENL-USPT-BRD4 axis in the brain microglia to HIV latency reversal. Brain tissue samples were obtained from the Last Gift cohort, including uninfected individuals, viremic, and PWH receiving suppressive ART to profile and integrate CNS cellular transcriptomes. (A) UMAP showed cells expressing BRD4 (red), MLLT1 (teal), and USP7 (yellow) in the different cell types. (B) Dot plot expressions of BRD4, MLLT1, and USP7 across different conditions (uninfected, viremic, ART-suppressed) were shown. Expression of BRD4, MLLT1, and USP7 in viremic (red) and ART-suppressed (yellow) donors compared to uninfected (grey) (C). The number of MLLT1+ (D), BRD4+ (E), and USP7+ cells (F) was shown in different cell populations.

**Figure S5.**
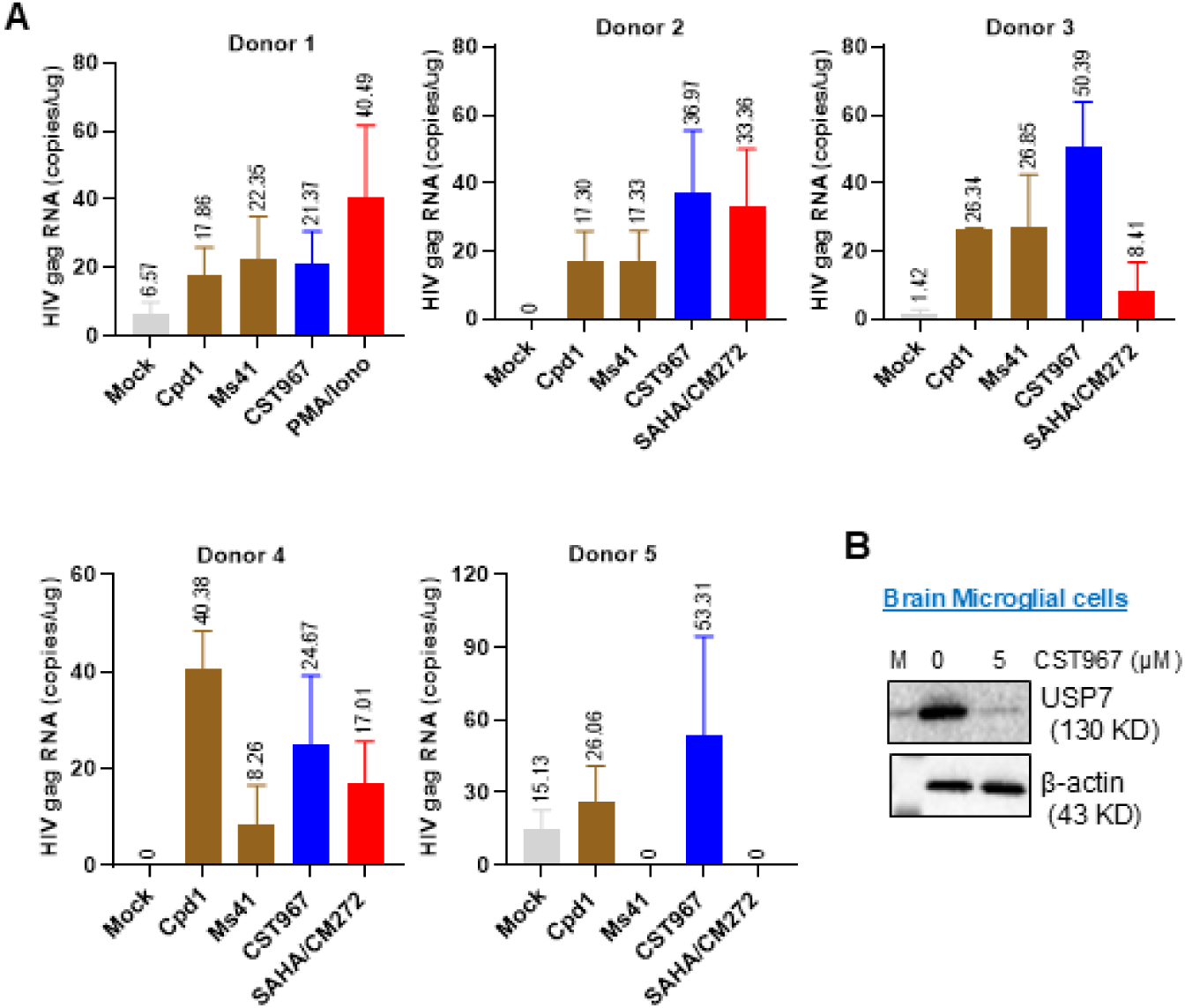
Targeting ENL-USP7 induces HIV latency reversal in brain microglia from PWH receiving suppressive ART. Brain microglia were isolated from five PWH receiving suppressive ART and treated with Cpd-1 (500 or 1,000 nM), MS41 (1,000 nM), CST967 (5 µM), PMA plus ionomycin (200 ng/mL and 2 µM), the HDAC inhibitors SAHA (250 nM) or CM272 (50 nM), or DMSO for 24 h. The data represent responses from individual donors (A). Cell-associated HIV *gag* levels were quantified by ddPCR, and results are shown for each donor with triplicate technical replicates. USP7 degradation was verified by immunoblotting (B).

**Figure S6.**
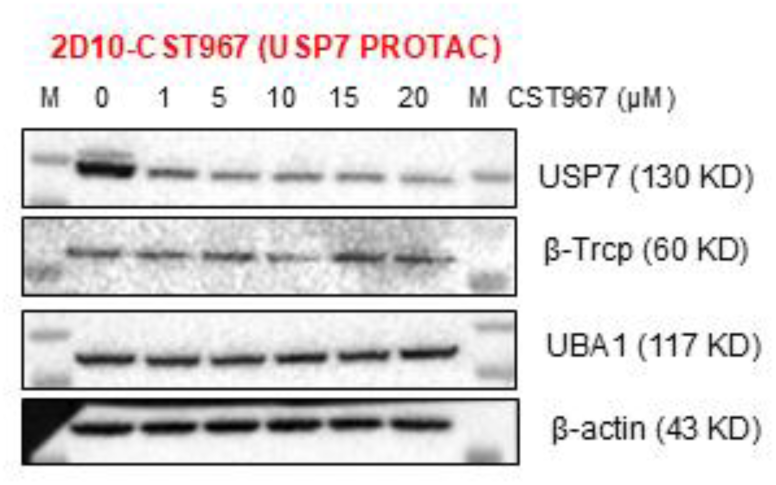
Examining the possible role of E3 ligases β-TrCP and UBA1 in ENL- and USP7-associated BRD4 instability. 2D10 cells were treated with the USP7 PROTAC CST967 for 24 h. Protein expression β-TrCP and UBA1was analyzed by western blot. β-Actin served as a loading control.

## Supplementary Tables

**Table S1.** Expression analysis of Protein ID from ENL proximity-biotinylation.

**Table S2.** Heat Map of SEC regulators identified in ENL pull down.

**Table S3.** Clinical information with PWH used for T cell latency reversal.

**Table S4.** Clinical demographics of the “Last Gift” cohort.

**Table S5.** Reagent or Resources used.

