## Supplementary figures and images for "The ENL–USP7 Complex Regulates HIV Latency Through BRD4 Stabilization"

### Table S3.docx

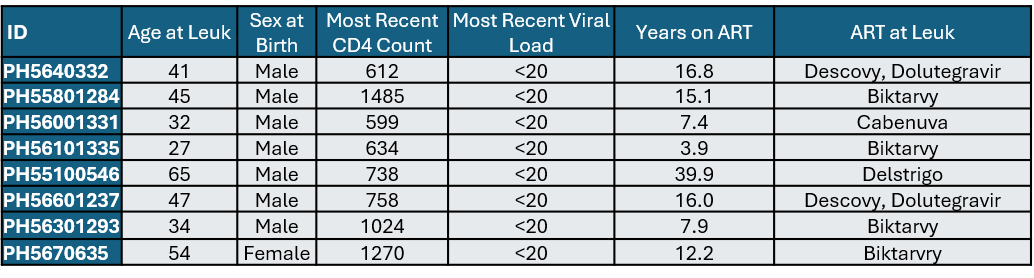

**Table S3. Clinical information with PWH used for T cell latency reversal.**

### Table S4.docx

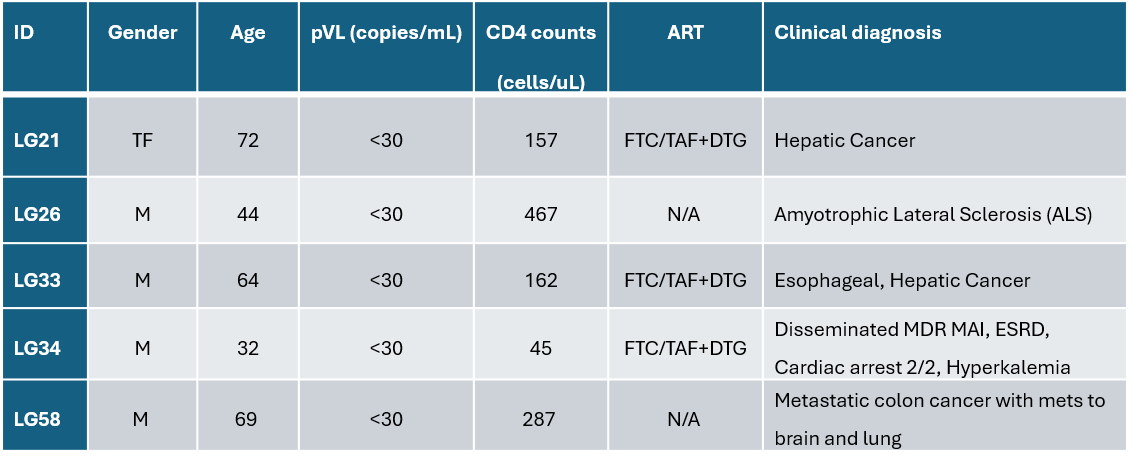

**Table S4. Clinical demographics of the “Last Gift” cohort.**
