## Supplementary Table 5 for "The ENL–USP7 Complex Regulates HIV Latency Through BRD4 Stabilization": Table S5.docx

**Table S5. Key Resources**

| REAGENT or RESOURCE | SOURCE | IDENTIFIER |
| --- | --- | --- |
| Antibodies |  |  |
| anti-ENL | Cell Signaling | Cat# 14893 |
| Anti-β-Actin | Cell Signaling | Cat# 4970 |
| anti-USP7 | Bethyl | Cat# A300-033A |
| Anti-Cyclin T1 | Cell Signaling | Cat# 81464T |
| Anti-CDK9 | Cell Signaling | Cat# 2316T |
| Anti-BRD4 | Cell Signaling | Cat# 13440T |
| Anti-BRD4 | Abcam | Cat# ab128874 |
| Anti-AF9 | Bethyl | Cat# A300-595A |
| Anti- β-Trcp | Cell Signaling | Cat# 11984 |
| Anti-UBA1 | Cell Signaling | Cat# 4890 |
| Anti-Flag-tag | Cell Signaling | Cat# 2368 |
| Anti-Myc-tag | Cell Signaling | Cat# 2278 |
| Normal rabbit IgG | Cell Signaling | Cat# 2729 |
| Anti-rabbit IgG, HRP-linked Antibody | Cell Signaling | Cat# 7074 |
| Anti-Streptavidin-HRP | Cell Signaling | Cat# 3999S |
| Biological samples |  |  |
| Leukapheresis from HIV-infected individuals | University of North Carolina Hospital | N/A |
| Chemicals, peptides, and recombinant proteins | | |
| Seradigm, Premium Grade FBS | Avantor | Cat# 97068-085 |
| Penicillin-Streptomycin | Gibco | Cat# 15140122 |
| L-Glutamine | Gibco | Cat# 25030081 |
| HEPES | Gibco | Cat# 15630106 |
| Sodium Pyruvate | Gibco | Cat# 11360070 |
| Recombinant Human IL-2 | PeproTech | Cat# 200-02 |
| anti-CD3/CD28 dynabeads | Gibco | Cat# 11132D |
| DNase I | Invitrogen | Cat# 18047019 |
| SuperScript™ III Reverse Transcriptase | Invitrogen | Cat# 18080093 |
| LIVE/DEAD™ Fixable Far Red Dead Cell Stain Kit | Invitrogen | Cat# L34973 |
| RIPA lysis buffer | Sigma-Aldrich | Cat# R0278 |
| Protease/Phosphatase Inhibitor Cocktail (100X) | Cell signaling | Cat# 5872 |
| NuPAGE™ LDS Sample Buffer (4X) | Invitrogen | Cat# NP0007 |
| Random primers | Invitrogen | Cat# 48190011 |
| Pierce™ 16% Formaldehyde (w/v), Methanol-free | Thermo Scientific™ | Cat# 28906 |
| PBS | Gibco | Cat# 14190-250 |
| NaCl (5 M), RNase-free | Invitrogen | Cat# AM9759 |
| IGEPAL® CA-630 (NP40) | Sigma-Aldrich | Cat# I8896 |
| Triton™ X-100 | Sigma-Aldrich | Cat# T8787 |
| Tris (1 M), pH 8.0, RNase-free | Invitrogen | Cat# AM9856 |
| EDTA (0.5 M), pH 8.0, RNase-free | Invitrogen | Cat# AM9260G |
| EGTA 0.5M, pH 8.0, Sterile | BioWORLD | Cat# 40520008 |
| SDS, 20% Solution | ThermoFisher | Cat# AM9820 |
| Nanodroplet cavitation reagent | MegaShear, Triangle Biotechnology | Cat# CS101-1000 |
| CST 967 (Usp7 PROTAC) | Tocris | Cat# 7801 |
| blasticidin | Medchemexpress | Cat# HY-K1054 |
| SR-0813 (ENL inhibitor) | Medchemexpress | Cat# HY-145409 |
| SAHA | Selleckchem chemicals | Cat# S1047 |
| Critical commercial assays |  |  |
| RNeasy mini kit | Qiagen | Cat# 74106 |
| TaqMan™ Universal PCR Master Mix | Applied Biosystems Inc. | Cat# 4304437 |
| Lipofectamine™ 3000 Transfection Reagent | Invitrogen | Cat# L3000015 |
| SYBR™ Green Universal Master Mix | Applied Biosystems Inc. | Cat# 4309155 |
| EasySep Release Human CD3 Positive Selection Kit | STEMCELL Technologies | Cat# 17751 |
| CD11b MicroBeads, human and mouse | Miltenyi Biotec | Cat# 130-049-601 |
| NE-PER™ Nuclear and Cytoplasmic Extraction Reagent | Thermo Scientific | Cat#78835 |
| Pierce™ Protein A/G Agarose | Thermo Scientific | Cat#20421 |
| Custom EasySep™ Human Resting CD4+ T Cell Isolation Kit | Stemcell Tech. | N/A |
| Pierce™ BCA Protein Assay Kits | Thermo Scientific | Cat# 23227 |
| Bio-Safe Coomassie Premixed Staining Solution | Biorad | Cat# 1610786 |
| ChIP Clean & Concentration kit | Zymo Research | Cat# D5205 |
| Experimental models: Cell lines |  |  |
| 2D10 | Dr. Jonathan Karn | N/A |
| J-Lat A1 | NIH HIV Reagent Program | N/A |
| J-Lat 10.6 | NIH HIV Reagent Program | N/A |
| U1 | NIH HIV Reagent Program | N/A |
| Oligonucleotides |  |  |
| HIV gag probe: FAM/CT CTC TCC T/ZEN/T CTA GCC  TCC GCT AGT /3IABkFQ/ | IDT | Malnati et al., 2008 |
| HIV gag FWD: TACTGACGCTCTCGCACC | IDT | Malnati et al., 2008 |
| HIV gag REV: TCTCGACGCAGGACTCG | IDT | Malnati et al., 2008 |
| SDHA (Hs00188166_m1) | Thermo Scientific | Cat# 4331182 |
| USP7 probe (Hs00931763_m1) | Thermo Scientific | Cat#4331182 |
| ENL probe (Hs00172962_m1) | Thermo Scientific | Cat#4331182 |
| BRD4 Probe (Hs04188087_m1) | Thermo Scientific | Cat#4331182 |
| ENL gRNA AA: CGGAGGGGTTCACTCACGAC TGG | Integrated DNA Technologies | Cat# Hs.Cas9.MLLT1.1.AA |
| ENL gRNA AB: CGACTCCTCTACTTTGTAGG GGG | Integrated DNA Technologies | Cat# Hs.Cas9.MLLT1.1.AB |
| ENL gRNA AC: GTTCCGGTACAAGCTCCTGC GGG | Integrated DNA Technologies | Cat# Hs.Cas9.MLLT1.1.AC |
| USP7 gRNA AA: CTACGTCGGCTTAAAGAATC AGG | Integrated DNA Technologies | Cat Hs.Cas9.USP7.1.AA |
| USP7 gRNA AB: TGATGGACACAACACCGCGG AGG | Integrated DNA Technologies | Cat Hs.Cas9.USP7.1.AB |
| USP7 gRNA AC: GTGTACATGATGCCAACCGA GGG | Integrated DNA Technologies | Cat Hs.Cas9.USP7.1.AC |
| Plasmid |  |  |
| pcDNA3.1-N-Myc_WT USP7 | Addgene | Plasmid# 131242 |
| pcDNA3.1-N-Myc_C223S USP7 | Addgene | Plasmid# 131243 |
| pcDNA5-Flag-BRD4-WT | Addgene | Plasmid# 90331 |
| pXLone-HA/Tet3G-3xFLAG/miniTurbo:Linker:hMLLT1 | VectorBuilder | N/A |
| Software and algorithms |  |  |
| FlowJo V10 | BD | https://www.flowjo.com/ |
| GraphPad Prism 10 | GraphPad Software Inc. | https://www.graphpad.com/ |
| SRplot |  | https://www.bioinformatics.com.cn/srplot |
| Cytoscape |  | https://cytoscape.org/ |
| ShinyGO 0.85.1 |  | https://bioinformatics.sdstate.edu/go/ |
| Biorander | Biorander | https://www.biorender.com/ |
| Cell Ranger v4.0 | R studio | 10x Genomics |
| Trailmaker | Parse Biosciences | https://app.trailmaker.parsebiosciences.com/ |
| Seurat v4.3.0 | R studio | https://github.com/satijalab/seurat |
